# Fast Alignment-Free Similarity Estimation By Tensor Sketching

**DOI:** 10.1101/2020.11.13.381814

**Authors:** Amir Joudaki, Gunnar Rätsch, André Kahles

## Abstract

The sharp increase in next-generation sequencing technologies’ capacity has created a demand for algorithms capable of quickly searching a large corpus of biological sequences. The complexity of biological variability and the magnitude of existing data sets have impeded finding algorithms with guaranteed accuracy that efficiently run in practice. Our main contribution is the Tensor Sketch method that efficiently and accurately estimates edit distances. In our experiments, Tensor Sketch had 0.956 Spearman’s rank correlation with the exact edit distance, improving its best competitor Ordered MinHash by 23%, while running almost 5 times faster. Finally, all sketches can be updated dynamically if the input is a sequence stream, making it appealing for large-scale applications where data cannot fit into memory.

Conceptually, our approach has three steps: 1) represent sequences as tensors over their sub-sequences, 2) apply tensor sketching that preserves tensor inner products, 3) implicitly compute the sketch. The sub-sequences, which are not necessarily contiguous pieces of the sequence, allow us to outperform *k*-mer-based methods, such as min-hash sketching over a set of *k*-mers. Typically, the number of sub-sequences grows exponentially with the sub-sequence length, introducing both memory and time overheads. We directly address this problem in steps 2 and 3 of our method. While the sketching of rank-1 or super-symmetric tensors is known to admit efficient sketching, the sub-sequence tensor does not satisfy either of these properties. Hence, we propose a new sketching scheme that completely avoids the need for constructing the ambient space.

Our tensor-sketching technique’s main advantages are three-fold: 1) Tensor Sketch has higher accuracy than any of the other assessed sketching methods used in practice. 2) All sketches can be computed in a streaming fashion, leading to significant time and memory savings when there is overlap between input sequences. 3) It is straightforward to extend tensor sketching to different settings leading to efficient methods for related sequence analysis tasks. We view tensor sketching as a framework to tackle a wide range of relevant bioinformatics problems, and we are confident that it can bring significant improvements for applications based on edit distance estimation.

## 1 Introduction

The emergence of next-generation sequencing technologies and a dramatic decrease in cost have led to an exponential increase in biological sequence data, frequently stored in exceptionally large databases. While alignment scores are considered the gold-standard of sequence distance in many bioinformatics applications, the growing number of sequences to be analyzed poses serious challenges to exact distance computation via alignment. This problem has motivated research on time- and space-efficient alignment-free methods that try to estimate rather than compute sequence similarity. Especially applications relying on comparing millions of sequences have turned towards choosing approximate methods over exact alignments [2]. Out of many possibilities, we have selected the task of phylogeny reconstruction to further motivate distance estimation. In this task, the estimated distances between sequences are used to reconstruct the evolutionary history in form of a phylogenetic tree. Instead of using exact alignment scores, many alignment-free methods instead rely on *k*-mer statistics as a proxy. (Where the term *k-mer* refers to all length *k* substrings of a given string.) The multiplicity, frequency, mode, and reappearance statistics of *k*-mers have been utilised to directly estimate evolutionary distance [17,16,1]. Other approaches include variable lengths matches in their statistics [18].

To break the stringent dependency of adjacent letters in a *k*-mer, *spaced k-mers* have introduced “match” and “ignore” positions. For example, if a match-pattern “<u>11</u>0<u>11</u>” is used, then “<u>CT</u>G<u>AC</u>” versus “<u>CT</u>T<u>AC</u>” constitutes a match. Spaced *k*-mers have been shown to improve mapping sensitivity, the accuracy of phylogenies, and the performance of sequence classification [5,7,13,15]. Analogously, substring-based methods can be relaxed to allow for some mismatches [6]. Since the quality of the estimations will greatly depend on the selected match-pattern, several works have focused on optimizing these patterns [3,11,8,10]. However, finding combinatorially optimal patterns becomes intractable as the number of ignore-positions increases. Furthermore, any optimization is prone to be task-dependent, which would require the optimization to be repeated for each task separately.

More recently, hashing based methods have become increasingly popular. MinHash sketch methods have been primarily used for set similarity estimation [2], that have shown promise in fast genomic and metagenomic distance estimation by representing sequences as collections of informative substrings [12]. From locality sensitive hashing literature we know that any sketch over the *ℓ^p^*-norm (*p* ∈ (0, 2]) will automatically lead to sub-quadratic nearest neighbor search by locality sensitive hashing [4]. This provides further motivation for focusing on sketching the edit distance, leaving behind the inherent problems of seed-based methods. One of the most accurate sketching methods currently available is *Ordered MinHash (OMH)* [9], which considers tuples *k*-mers that are non-adjacent. Therefore, we will compare our approach against the classical *k*-mer methods MinHash, and Ordered MinHash.

An especially difficult task for most current methods is the estimation of edit distance for longer, less closely related sequences. To illustrate the main limitations of using *k*-mer statistics in this context, we look at two examples that act as opposing forces in deciding the size of *k*.

### Example 1.

Consider two sequences 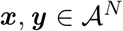 drawn randomly over the alphabet of size 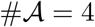, and let *v_x_, v_y_* ∈ ℝ^4^*k*^^ denote the *k*-mer frequency profile (multiplicity of each *k*-mer divided by total number of k-mers) for each sequence. For long sequences with *N* ≫ 4^*k*^, *k*-mer frequencies will converge to their mean, that is 4^-*k*^ for all their components, which implies that ||*v_x_* — *v_y_* ||_1_ → 0. Therefore, any *k*-mer profile-based method will severely *underestimate* distance between these two random sequences. In order to avoid this, k has to be restricted to larger values *k* ≳ log_4_(*N*).

### Example 2.

Consider the scenario that we want to rely on *k*-mer matching to find similar parts of two sequences ***y*** and ***x***, and ***y*** is generated by independently mutating every index of ***x*** with probability *r* ∈ (0,1). Therefore, the probability that a *k*-mer belonging to ***x*** and ***y*** is not mutated is 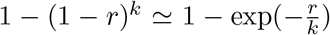. This implies that *k* ≲ *r*^-1^ is necessary to avoid the *k*-mer match probability to converge to zero.

By combining the insights from these examples, we conclude that any value of k will fail with high probability in one of these scenarios for distantly related sequences. Since both results hold for *k*-mer frequency vectors in general, any such statistic on the *k*-mer profile is also bound to fail. This inherent problem with k-mers is our main motivation for designing sketches that are more resilient to mutations.

Conceptually, our method can be seen as expectation over all possible spaced seeds, without any limits on the number of *ignore* positions. In other words, Tensor Sketching resolves the inherent limitations of any seed-based method by taking an average of all possible seeds, and thereby reducing the risk of selecting a seed that reduces the sensitivity. In statistical terms, this implies that the resulting sketch is a Bayesian admissible estimator.

We will begin by introducing our notation and terminology in Section 2.1 and then proceed to presenting the Tensor Sketch and Tensor Slide sketch methods in Sections 2.2 and 2.3, respectively. In subsequent Section 3 we first summarize our data generation scheme and then compare our approach to the available state-of-the-art methods. We then motivate possible applications such as phylogeny reconstruction and graph distance estimation. In the final Section 4 we put our work into context and give an outlook into future work.

## 2 Methods

### 2.1 Preliminaries and Notation

#### Sets

We use 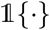 to denote the indicator function, taking a value of 1 when the logical expression between the brackets is true, and zero otherwise. For integer *N*, [*N*] denotes the integer set [*N*]: = {1,…, *N*}. For finite set *S*, let |*S*| and #*S* interchangeably denote its cardinality, and define to be all bijective maps from *S* to [|*S*|]

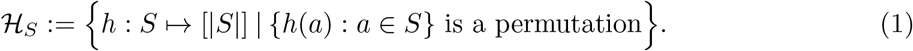

Finally, we use *X* ~ *S* to denote that *X* is uniformly drawn from the set *S*.

#### Vectors

Throughout the manuscript bold face fonts are used to distinguish vectors, tensors, and sequences, from scalar variables **a** = (*a*_1_, *a*_2_). **1**^*N*^ and **0**^*N*^ are all-ones and all-zeros vectors of *N* length *N*, respectively. We use 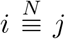 when integers *i* and *j* are equal modulus *N*. The circular *r*-shift shift_*r*_ (**a**), will shift the elements of **a** to the left shift_*r*_ (**a**) = (*a*_*r*+1_,… *a_N_, a*_1_,…,*a_r_*), defined formally as

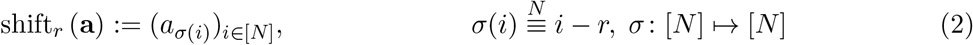

where the mapping *σ*, circularly shifts indices to the left.

#### Strings

Let 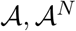, and 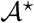 denote the alphabet, the strings of length *N*, and all strings respectively. Let ***x*** o ***y*** denote concatenation of ***x*** and ***y***, and |***x***| be the length of string ***x***. We use ***x***_*i*_ to denote the *i*-th character. Define ***x***_[*i:j*]_ to be a slice from ith to *j* th index ***x***_[*i:j*]_: = ***x***_*i*_… ***x**_j_*, referred to as a *substring* of ***x***, and define ***x***:= ***x***_*i*_1__ … ***x***_*i_k_*_ to be a *subsequence* of ***x***, when 1 ≤ *i*_1_ < · · · < *i_k_* ≤ |***x***| are strictly increasing indices of ***x***. For two strings 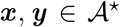, the edit distance d_ed_(***x, y***) denotes the minimum number of edit operations needed to transform one string to the other. It can be defined recursively using 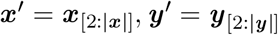 as follows

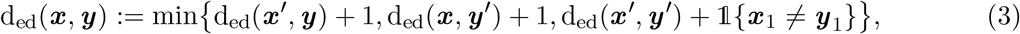

with d_ed_(*ε,ε*): = 0 as the recursion basis.

#### Minimum hash sketching

For a sequence 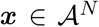, let 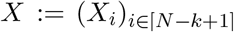 denote its *k*-mer sequence, with *X_i_* denoting the ith *k*-mer *X_i_*:= ***x***[_*i:i*+*k*-1_], and #*X_i_* the occurrence number of this *k*-mer #*X_i_*:= #{*j ≤ i*: *X_j_* = *X_i_*}. Furthermore, the pairs (*X_i_*, # *X_i_*) will be referred to as *unified k-mers*.

In *MinHash (MH),* a random hash is used to sort the *k*-mers, the *k*-mer with the lowest index, with respect to hash functions 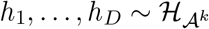

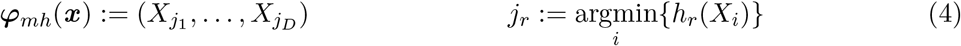

In *Weighted MinHash (WMH),* hash functions are drawn from a hash family over unified *k*-mers 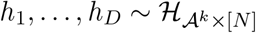

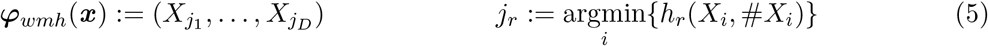

Finally, *Ordered MinHash (OMH)* [9], generalizes WMH by sampling from multiple *k*-mers that appear in the same order within the sequence. The sampling uses hashes over the *k*-mers space 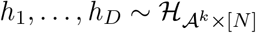, and selects indices that are mapped to the t lowest hash values. Formally, for each hash function *h_r_*, we sketch (*X*_*σ*_1__,…,*X_σ_t__*), when monotone indices 1 ≤ *σ*_1_ < · · · < *σ*\* ≤ |*X*| are mapped to the lowest hash values

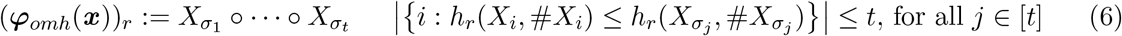

We can compute distances via the *Hamming distance* d_H_(*a, b*), defined as the number of indices that the input sequences differ d_H_(*a, b*) = #{*i*: *a_i_* = *b_i_*}, where *a* and *b* have equal length. When *φ* is a MinHash sketch, d_H_(*φ*(***x***),*φ*(***y***)) is closely related to the Jaccard set similarity index between *k*-mer sets

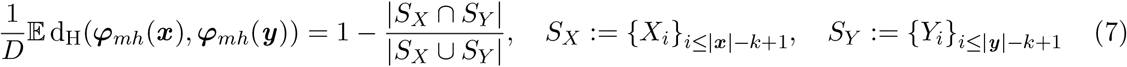

Note that MH, WMH, and OMH are defined as elements with lowest hash over a set of *k*-mers, unified *k*-mers, and ordered tuples of *k*-mers, respectively. For example, in WMH we can change the definition of *S_X_* to a set of unified *k*-mers *S_X_*:= {(*X_i_*, #*X_i_*): *i* ≤ |***x***| — *k* + 1} (analogous for *S_Y_*). It is worth mentioning that OMH becomes equivalent to WMH if we set the tuple length to one, *t* = 1.

### 2.2 Tensor Sketch

First, define 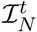 to be set of all increasing subsequences of [*N*]*

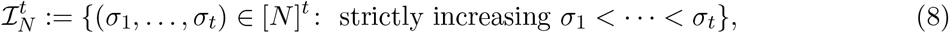

Without loss of generality let the alphabet be relabeled to 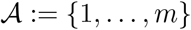, and 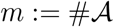 be the size. The main idea behind sequence tensor sketch is counting subsequences as opposed to *k*-mers. Define 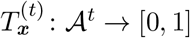 to be the probability mass function of a random *t*-ary subsequence of *x*

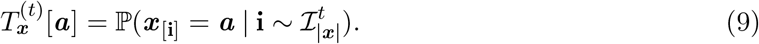

With a slight abuse of notation, we treat 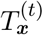 both as a probability mass function, and a *t*-dimensional tensor with size *m^t^*. Then, for two arbitrary strings 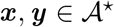, define order t similarity *s*^(*t*)^(***x, y***) and distance *d*^(*t*)^(***x, y***) as

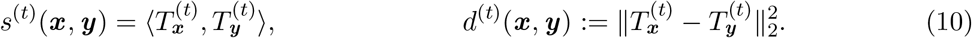

The inner product equals the probability of getting a match, if we draw two *t*-ary tuples from ***x*** and ***y***, respectively, which is closely related to OMH sketch.

Naive computation of equations (9) and (10) requires 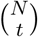 time and *m^t^* memory, which becomes prohibitive for even moderate values of these variables. Our tensor sketching scheme provides a (1+*ϵ*)-approximation for tensor distance, but only requires 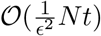 time to compute. Furthermore, any downstream analysis will operate on sketches of size 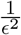, regardless of the original sequence length.

Since we are interested in preserving the Euclidean norm after sketching, we follow the definition of tensor sketch by Pham and Pagh [14]. For integer *D*, tensor sketch *Φ*: ℝ^*m*^*t*^^ → ℝ^*D*^ sketches an *m^t^*-dimensional tensor into ℝ^*D*^. Let us assume a series of pairwise independent hash functions 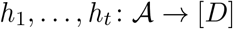, and sign hash functions 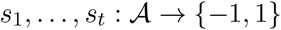. Moreover, define hash sum 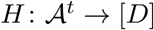 and hash sign product 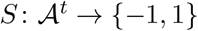 as follows

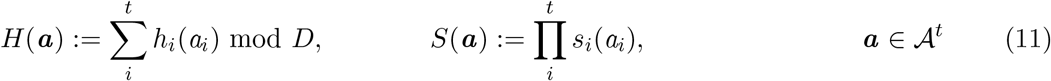

Finally, tensor sketch Φ(*T*): = (*ϕ_r_* (*T*))_*r*∈[*D*]_ is defined as

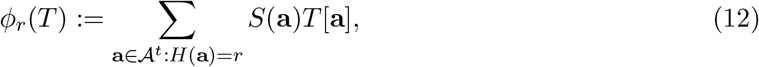

where *T* ∈ ℝ*^m^t^^* is an arbitrary tensor. Crucially, tensor sketch preserves the Euclidean norm in expectation 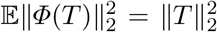, and assume t to be constant, the variance decreases with sketch dimension 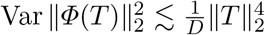. Therefore, 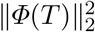 is concentrated around its mean by second moment bound (see Lemma 7 of Pham and Pagh [14]). Therefore, a sketch size of 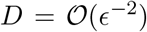 suffices to bound the multiplicative error by *ϵ*.

If we could compute 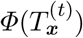 and 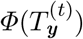) efficiently, distance computation 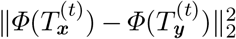 merely requires 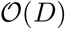 time, as opposed to the exponential cost of constructing the ambient space. However, since the tensors 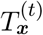 are never rank-1, the conventional efficient tensor sketching schemes cannot be applied here. Therefore, we have designed an efficient algorithm to compute (12), and provide several extensions of the algorithm in the next section. In summary, for sequence ***x***, the tensor sketch ***φ_ts_***(***x***) is computed by applying the tensor sketch on tuple tensor 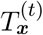

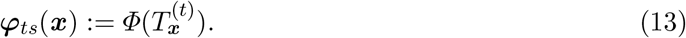

#### Implicit sketching by recursive computation

First, observe that the rth component of the sketch 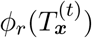, can be rewritten based on the probability mass of (*H*(***x***_[**i**]_),*S*(***x***_[**i**]_)), when **i** is a uniformly drawn ordered *t*-tuple 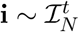

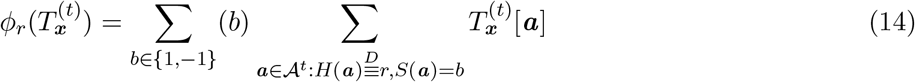

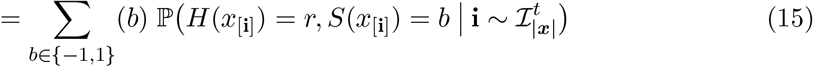

We set out to recursively compute the probability mass for t-ary tuples, based on probability mass of shorter ordered tuples. Therefore, for *p* ∈ {1,…, *t*} let 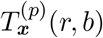 be the probability mass function of 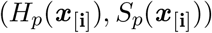, when 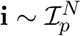 is a random ordered p-tuple over [*N*], while 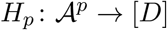 and 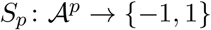 are partial hash sum and products up to *p*, respectively

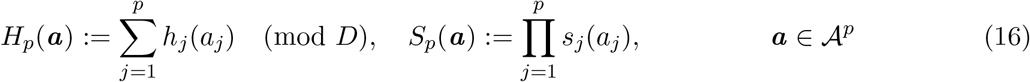

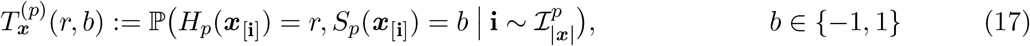

To recursively compute 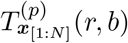 for *p* ≥ 2, we separate it into two cases depending on whether the last index is part of the tuple (*i_p_* = *N*) or not (*i_p_* < *N*)

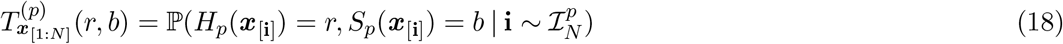

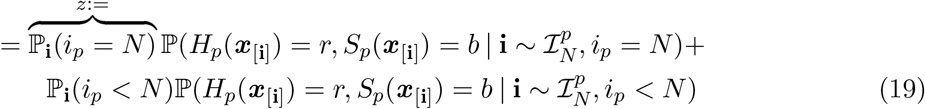

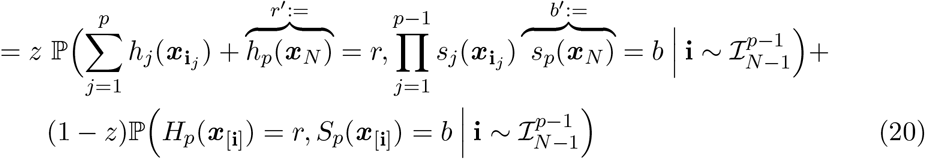

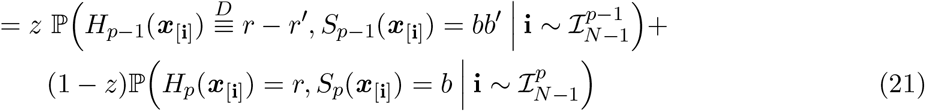

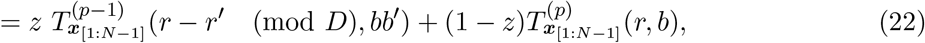

where 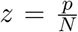, since 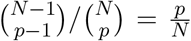. Defining vector 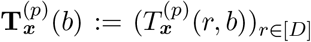, the index shift *r – r*’ (mod *D*), amounts to circularly shifting of 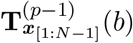 to the left by *r*’ indices. Hence, the recursion in vector form can be written as

♢*Recursive insert* (*p* ∈ [*t*]):

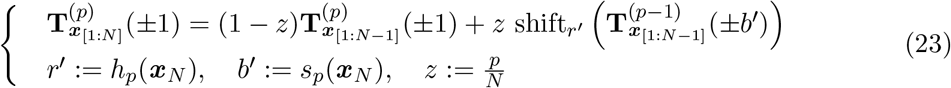

♢ *Basis* (*p* = 0):

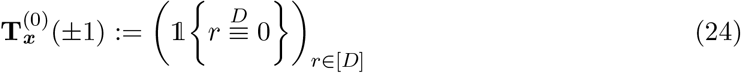

Interestingly, the recursion relation for variables in 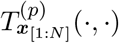 depends only on variables in the previous layer 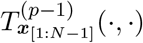. Therefore, it is sufficient to store the variables corresponding to layer *i*, which implies a memory complexity of 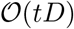. This fact is exploited in Algorithm 1, which is why the subscript ***x***_[**i**]_ is dropped from **T**, and uses ±1 for brevity^4^. The number of sub-problems needed to compute 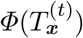 is exactly 2*NtD*, and the cost of each recursion is 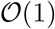. Therefore, we can compute the sketch in 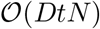 time.

**Algorithm 1:**
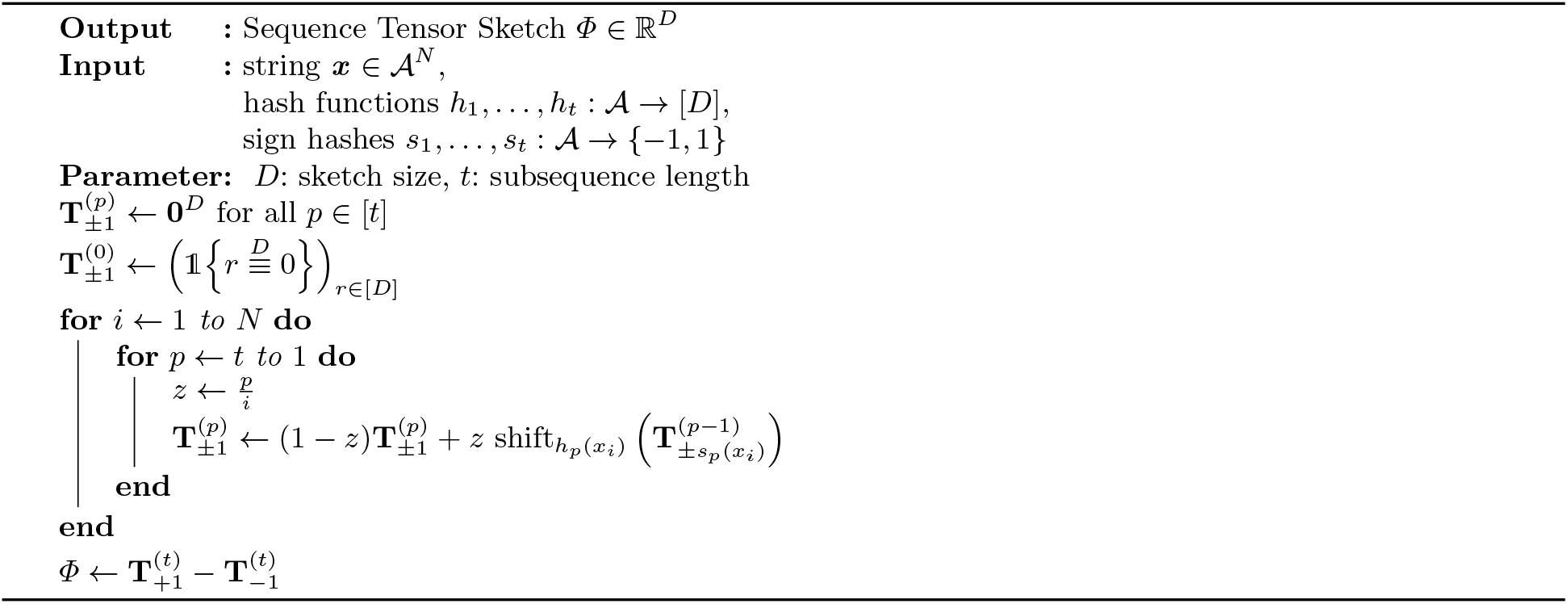
Sequence Tensor Sketching

### 2.3 Tensor Slide Sketch

In this section, we improve the precision of tensor sketch by concatenating sketches of overlapping subsequences of the original sequence. However, instead of applying Algorithm 1 to compute each sketch separately, we can exploit the fact that many sub-problems in the recursion (18) are shared. The main idea is to consider sub-problems 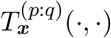, that correspond to hash functions *s_p_,…,s_q_* and *h_p_,…,h_q_*, for all 1 ≤ *p* ≤ *q* ≤ *t*, so that we can efficiently remove characters from the initial position. Formally, 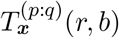 for all *r* ∈ [*D*],*b* ∈ {—1,1}, is the probability of 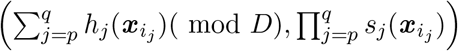 being equal to (*r, b*), when **i** is a randomly drawn (*q – p*+1)- ary increasing subsequence 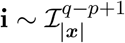 (See **Supplementary Section** A for more details).

In Algorithm 2, 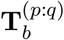 denotes the probability vector 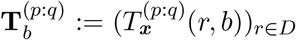. The time and memory complexity of computing sketches for all *w*-mers of a reference sequence with length *N* are 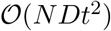 and 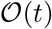, respectively. The length of the sliding window *w* does not appear in any complexity term.

Equipped with the streaming algorithm, we can concatenate sketches of windows as we slide the window along the sequence. More specifically, because sliding windows that have a large overlap cannot improve our estimate, we can down-sample by a factor of *S*, referred to as the stride. In other words, we store the sketch of a window every *s* basepairs. Formally, let us define *φ_tss_*(***x***) as follows

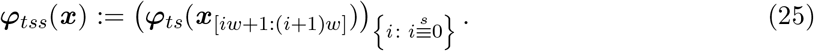

where *φ_ts_*(***x***) is Tensor Sketch(13). If we naively call tensor sketch for each step, 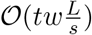 time is required per embedding dimension. Applying the tensor slide sketch algorithm, this can be computed in 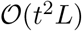. In our experience, moderate values of *t* ≤ 5 suffice for most applications, implying that for all 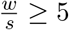, the streaming algorithm will run faster. Finally, the distance between two tensor slide sketches is defined as the squared Euclidean norm, with zero-padding for shorter sequences if two sequences are not equal in length.

**Algorithm 2:**
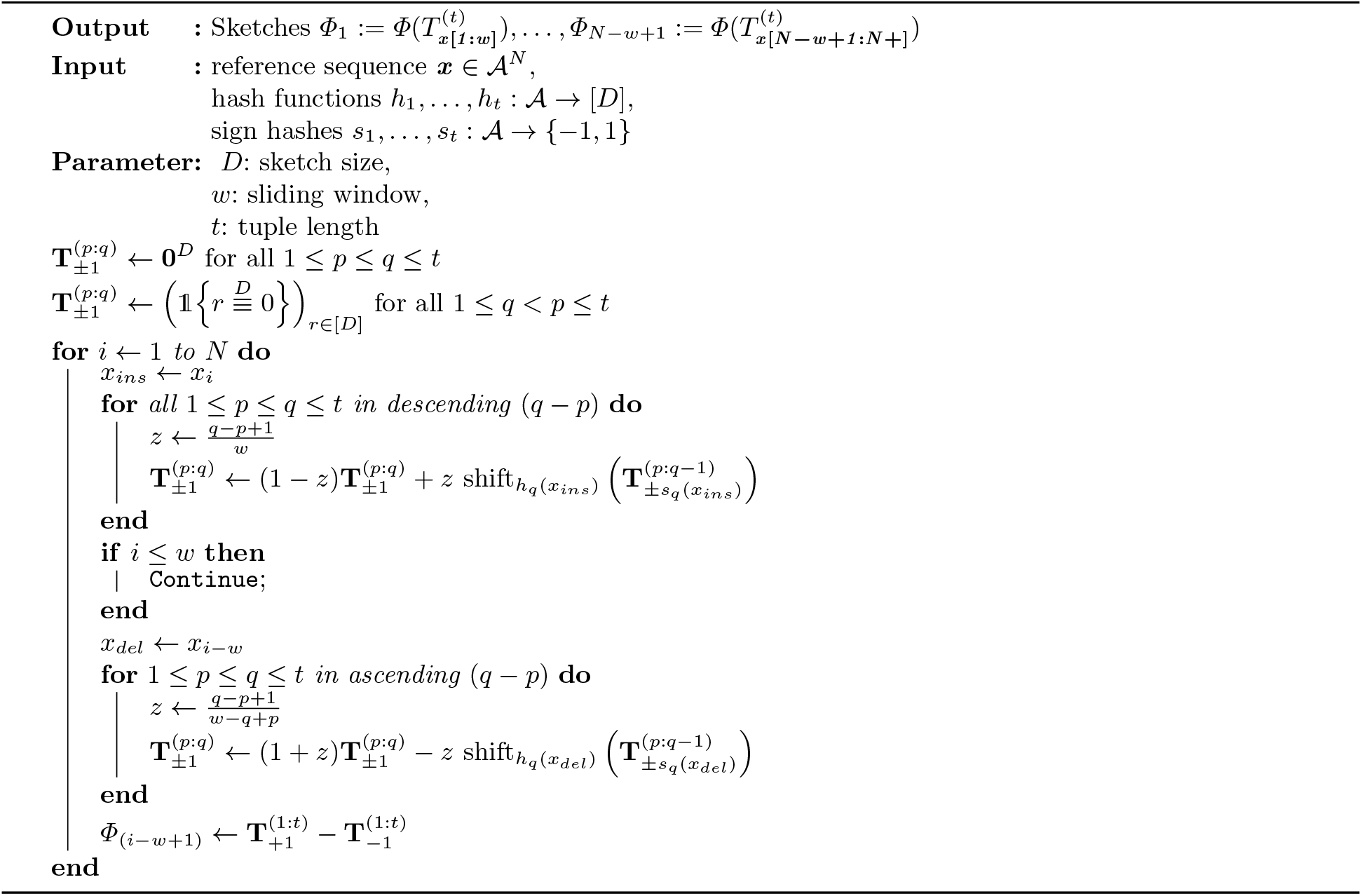
Tensor Slide Sketch3 Experimental Results

## 3. Experimental Results

The primary objective of our tensor sketching framework is to efficiently sketch global alignment scores between sequences. In order to put our method into context, we compare its performance to MH, WMH, and OMH [9], three state-of-the-art sketching techniques (see **Section** 2 for formal definitions).

### 3.1 Synthetic data generation and parameter selection

As a basis for evaluation, we generated a set of synthetic sequence pairs. Given a mutation rate *r* ∈ (0,1), and a reference sequence at random 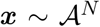, we mutate each index with probability *r*, with mutations being insertion, deletion, or substitution, with equal probabilities. While the mutation rate for each pair is fixed, it is uniformly drawn from the unit interval for each pair anew, which ensures the diversity of the edit distances.

One can evaluate the sketch-based distance of each pair against the exact edit distance, by quantifying how well each method preserves the order of distances, as captured by the Spearman’s correlation. We select parameters for each hashing method such that it achieves the highest rank correlation for a fixed embedding dimension *D* (See **Supplementary Section** B for more details regarding parameter selection).

### 3.2 Tensor slide sketch achieves high rank correlation with edit distance

For each sketching method, we determined Spearman’s rank correlation between edit distance and sketch distance, as well as absolute and relative execution time (**Table 1**). In particular, Tensor Slide Sketch achieves a rank correlation of 0.956 while reducing the computation time by 96.3%. Furthermore, we calculated the area under the receiver operating characteristic (AUROC) for a classifier discriminating two sequences based on a threshold for their exact edit distance (normalized by length), indicating that TS and TSS outperform other methods in detection of distantly related sequences (normalized edit distance threshold 0.5), while TSS outperforms other methods on all but the smallest edit distance threshold.

**Table 1:**
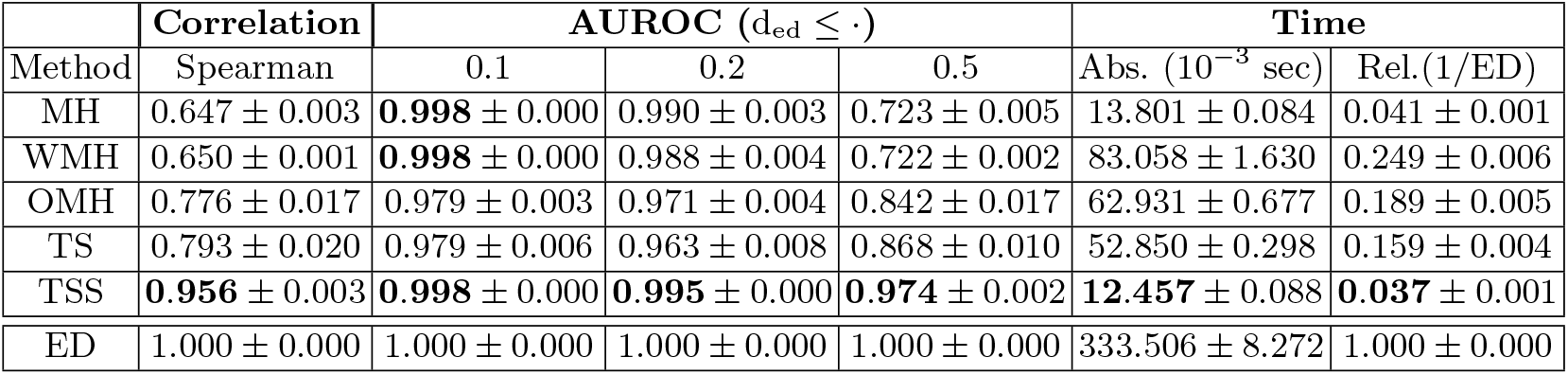
1000 sequence pairs of length 10000 were generated over an alphabet of size 4, with the mutation rate uniformly drawn from [0.00, 1.00]. The time column shows normalized time in milliseconds, i.e., total time divided by number of sequences, while the relative time shows the ratio of sketch-based time to the time for computing exact edit distance. The values shown are average over independent trials followed by their standard deviation. The embedding dimension is set to *D* = 64, and individual model parameters are (a) MinHash *k* = 12, (b) Weighted MinHash *k* = 12, (c) Ordered MinHash *k* = 2,*t* = 7, (d) Tensor Sketch *t* = 6, (e) Tensor Slide Sketch *w* = 1000, *s* = 100, *t* = 3.

| Method | Correlation | AUROC ($d_{\text{ed}} \leq \cdot$) | | | Time | |
| --- | --- | --- | --- | --- | --- | --- |
| | Spearman | 0.1 | 0.2 | 0.5 | Abs. ($10^{-3}$ sec) | Rel.(1/ED) |
| MH | $0.647 \pm 0.003$ | **$0.998 \pm 0.000$** | $0.990 \pm 0.003$ | $0.723 \pm 0.005$ | $13.801 \pm 0.084$ | $0.041 \pm 0.001$ |
| WMH | $0.650 \pm 0.001$ | **$0.998 \pm 0.000$** | $0.988 \pm 0.004$ | $0.722 \pm 0.002$ | $83.058 \pm 1.630$ | $0.249 \pm 0.006$ |
| OMH | $0.776 \pm 0.017$ | $0.979 \pm 0.003$ | $0.971 \pm 0.004$ | $0.842 \pm 0.017$ | $62.931 \pm 0.677$ | $0.189 \pm 0.005$ |
| TS | $0.793 \pm 0.020$ | $0.979 \pm 0.006$ | $0.963 \pm 0.008$ | $0.868 \pm 0.010$ | $52.850 \pm 0.298$ | $0.159 \pm 0.004$ |
| TSS | **$0.956 \pm 0.003$** | **$0.998 \pm 0.000$** | **$0.995 \pm 0.000$** | **$0.974 \pm 0.002$** | **$12.457 \pm 0.088$** | **$0.037 \pm 0.001$** |
| ED | $1.000 \pm 0.000$ | $1.000 \pm 0.000$ | $1.000 \pm 0.000$ | $1.000 \pm 0.000$ | $333.506 \pm 8.272$ | $1.000 \pm 0.000$ |

We then assessed the performance and execution time of all considered methods utilizing the synthetically generated data set (Figure 1). We observe that the quality of both MinHash (MH) and Weighted MinHash (WMH) greatly diminishes as the sequences become more distant from one another (**Figure 1a**) and the quality of MH and WMH drops with increasing sequence length (**Figure 1b**). In both plots, the accuracy of OMH and the tensor sketch methods maintains a much higher level. Notably, the increased accuracy of TS and TSS comes only at a very small additional cost in time when compared with MH (**Figure 1c**), while both methods run much faster than OMH. Lastly, we assessed the relationship of sketching accuracy to the number of embedding dimensions for each method (**Figure 1d**). Remarkably, TSS achieves a rank correlation of 0.74% even when using only 10 embedding dimensions.

**Fig. 1:**
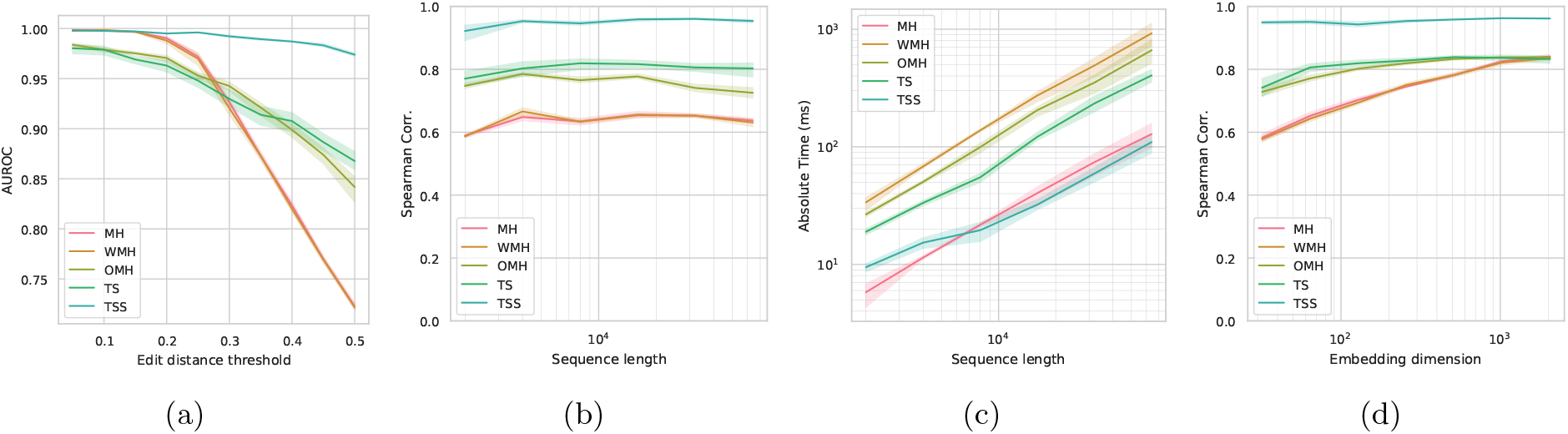
The dataset for these experiments consisted of 2000 sequence pairs independently generated over an alphabet of size 4. The embedding dimension is set to *D* = 256, and model-specific parameters are MinHash *k* = 12, Weighted MinHash *k* = 12, Ordered MinHash *k* = 2, *t* = 7, Tensor Sketch *t* = 6, Tensor Slide Sketch *t* = 3, with window and stride set to 10% and 1% of the sequence length, and its embedding dimension is set to the square root of D. (1a) Area Under the ROC Curve (AUROC), for detection of edit distances below a threshold using the sketch-based approximations. The x-axis, shows which edit distance (normalized) is used, and the y axis shows AUROC for various sketch-based distances. (1b) The Spearman’s rank correlation is plotted against the sequence length (logarithmic scale). (1c) Similar setting to (1b), plotting the execution time of each sketching method (y-axis, logarithmic scale) as a function of sequence length (x-axis, logarithmic scale). The reported times are normalized, i.e., average sketching time plus average distance computation time for each method. (1d) Spearman rank correlation of each sketching method as a function of the embedding dimension *D* (*x*-axis, logarithmic scale). In all subplots the solid line and shades indicate mean and standard deviation, computed over independent runs with the same configuration.

In summary, TSS produces the most accurate sketches across the board, while introducing a small time and memory overhead when compared with min-hashing. Furthermore, TS and TSS can be computed in a streaming fashion. For example, in order to sketch all 100-mers of a reference sequence, the time complexity grows only linearly with the length of the reference sequence, while WMH and MH need to be computed separately for each sequence of length m, rendering them *m* times slower for this task.

While sorting creates an iterative bottleneck for all hash-based schemes, the tensor sketch operation is based on a vector shift and sum, which explains the faster execution in practice. Moreover, vector operations can be executed in a single step on some CPU or GPU architectures to achieve an even greater boost. Finally, dependence on large values of k will introduce additional memory and time overhead if *k*-mers no longer fit into built-in types. In contrast, tensor sketch can be stored in efficient built-in types regardless of the choice of parameters.

### 3.3 Tensor Sketching helps estimate phylogenies

We further explored the task of estimating all pairwise distances of a given set of evolutionary related sequences, resulting in a well structured distance matrix. This task is reminiscent of important bioinformatics applications, such as the binning of metagenomic reads or phylogeny reconstruction. In both cases, the reconstruction of the good distance matrix as a whole, and not only individual components, forms the basis for a good approximation. While there are several ways to formulate this problem, we can simply visualize the exact and approximate matrices and compare them with an overall metric such as Spearman’s rank correlation. Figure 2, shows such a distance matrix. The sequences were generated to emulate a phylogeny tree. Furthermore, each sequence is mutated from its parent with a fixed rate of 15%. The optimal parameters of each model are chosen again based on the Spearman’s rank correlation, while window and stride length for TSS are fixed to 10% and 1% of the sequence length, giving an advantage to the competing methods. It is evident that MH and OMH are only accurate for small distances, but fail to do so for more distant sequences, which could negatively affect tree reconstruction based on these estimate distances. On the other hand, OMH and tensor sketching are capable of preserving a wider range of distances, which makes them better candidates for approximate phylogeny construction.

**Fig. 2:**
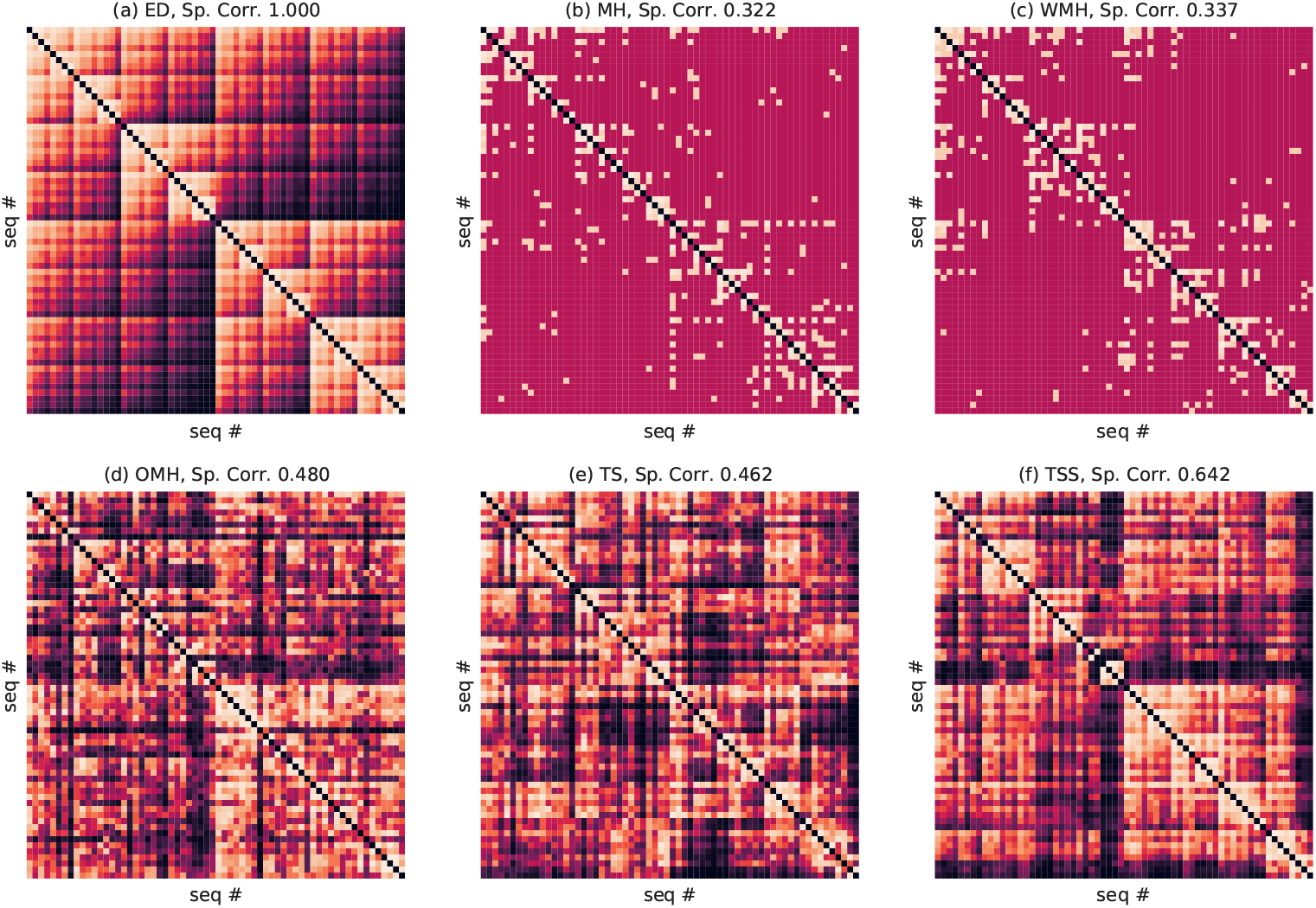
The subplot (a) illustrates the exact edit distance matrix, while the subplots (b)-(f) demonstrate the approximate distance matrices based on sketching methods. To highlight how well each method preserves the rank of distances, in all plots, the color-code indicates the rank of each distance (lighter, shorter distance), and the Spearman’s rank correlation is shown at the top of each plot. The phylogeny was simulated by an initial random sequence of length *N* = 10000, over an alphabet of size 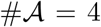. Afterwards, for 6 generations, each sequence in the gene pool was mutated and added back to the pool, resulting in 64 sequences overall. The mutation rate set to 0.15. The embedding dimension is set to *D* = 64, and individual model parameters are set to (b) MinHash *k* = 12, (c) Weighted MinHash *k* = 12, (d) Ordered MinHash *k* = 2, *t* = 7, (e) Tensor Sketch *t* = 5, (f) Tensor Slide Sketch *t* = 2, *w* = 1000, *s* = 100.

### 3.4 Tensor Sketch opens up a wide range of applications

#### Sketching de-Bruijn graph distances

The slide-sketch algorithm presented in 2, assumes only having access to a stream of characters coming in and that the task involves sketching substrings of the entire sequence. While the task description may sound to be restricted to a linear reference sequence, the same assumptions actually apply to any Euler path on a string graph. For example, if the input is a de-Bruijn graph and the task is to sketch all vertices of the graph, any traversal of the graph that walks through adjacent vertices can be mapped to the streaming of incoming and outgoing characters, which will fit the assumptions of the slide sketch algorithm. Consequently, the time complexity of sketching vertices of a de-Bruijn graph will grow only as a function of its size, i.e., the number of its vertices, as opposed to the size of each vertex.

#### Robustness to Global Mutations

Tensor Sketch is sensitive to global shifts and transpositions of characters, while *k*-mer statistics are entirely local. To illustrate this, consider case when sequence ***y*** is constructed from ***x***, by swapping the first and second half of the sequence 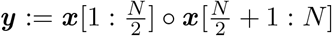. It is evident that (*N — k* + 1) out of *N k*-mers are intact under this block permutation, implying that for sufficiently large *N ≫ k* the *k*-mer profile of ***y*** will converge to ***x***. This may severely overestimate their alignment score. In contrast, only 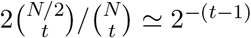 of *t*-ary subsequences remain intact. This greater sensitivity makes tensor sketch an appealing method for applications that such block permutations are likely to occur.

## 4 Discussion

We presented tensor sketching, a method for estimating sequence similarity for biological sequences. We demonstrated that Tensor Slide Sketch achieves a high Spearman’s rank correlation, but runs an order of magnitude faster than computing the exact alignment. When compared with other sketching methods, Tensor Sketch and Tensor Slide Sketch both achieve a more accurate preservation of the order between distances than MinHash, Weighted MinHash, and Ordered MinHash.

### Bayesian estimation with tensor sketching

Spaced *k*-mers were introduced motivated by applications involving distantly related sequences. If we are to allow for *i ignore* positions, there will be 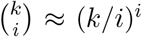 possible patterns. Considering all such patterns is clearly not an option if i is anything larger but a small constant. Not only is it highly non-trivial to search for the optimal pattern in this large combinatorial space, but any incremental step for pattern optimization also has to be repeated if different tasks require a different set of patterns. Seen from this angle, Tensor Sketch can be interpreted as an average of all possible spaced t-mers patterns with (*N — t*) ignore positions, while the sketching step avoids the exponential cost of explicitly representing all combinatorial options.

The core idea of tensor sketching, that is to average over all possible seed patterns, can be alternatively viewed as Bayesian estimation. This implies that our estimate is admissible, i.e., no other estimator can outperform it across the board. This provides some intuition into why our approach achieves an acceptable or better accuracy over the entire range of edit distances. In other words, the risk of selecting a bad seed or size of *k*-mer is minimized by taking the mean over all possible seeds. While this corresponds to a uniform prior on our seeds, one can introduce a non-uniform prior by setting different weights for vertices on our tensors, namely to penalize more number of gaps in the pattern. Therefore, we can strengthen or weaken the contribution of individual patterns. Weighted average can help to reduce background statistics into the distance metric, as well as introduce nonuniform mismatch penalties. These variations and extensions come naturally within the tensor sketching framework, in contrast with hash-based approaches that require heuristic steps.

### Conclusion

This work’s main contributions are the introduction of two tensor-based sketching methods to sequence similarity estimation: Tensor Sketch, providing an efficient algorithm to compute it, and Tensor Slide Sketch, a streaming version of the algorithm. Our results indicate that the tensor sketching method and its applications open up exciting research directions to explore for the bioinformatics community. The main advantages of tensor sketching compared with hash-based methods are that it 1) can run in a streaming fashion, 2) achieves much higher accuracy, and 3) runs much faster in practice.

## Acknowledgements

We would like to thank Ximena Bonilla, Daniel Danciu, Kjong-Van Lehmann, and Ragnar Groot Koerkamp for their constructive feedback on the manuscript. AJ was funded by ETH Zurich core funding to GR. AK was partially funded by PHRT Project #106 and the Swiss National Science Foundation Grant No. 407540_167331 “Scalable Genome Graph Data Structures for Metagenomics and Genome Annotation” as part of Swiss National Research Programme (NRP) 75 “Big Data”.

## Appendix A Tensor Slide Sketch Algorithm

For all 1 ≤ *p* ≤ *q* ≤ *t*, let 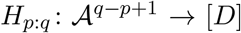 and 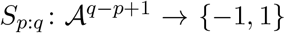 be partial hash sum and products from *p* up to *q*, respectively, and define 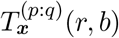 be the probability mass function of 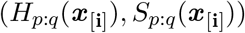, when **i** is a uniformly drawn ordered (*q — p* + 1)-tuple.

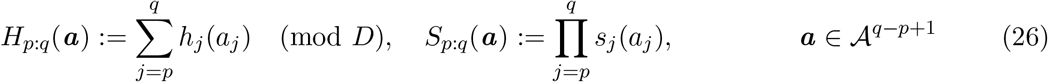

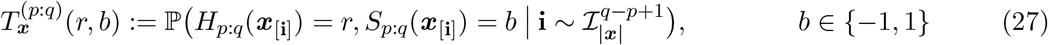

The recursive insertion is analogous to (18), as we can derive 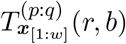 based on the previous layer 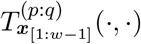 and smaller problems 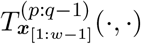. Recursive deletion is equivalent to rolling back insertion of the first character. We can derive 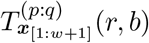 based on 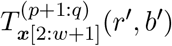 and 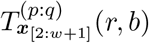, by separating it into two cases when the random tuple starts at the first index being in the tuple

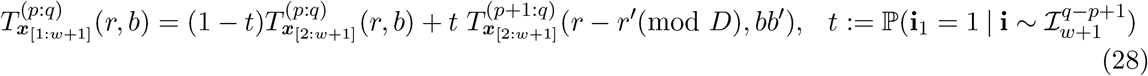

where 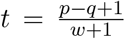, analogous to the insertion only case, and *r*’: = *h_p_*(***x***_1_) and *b*’: = *s_p_*(***x***_1_) are recursive hashes as before. We can rearrange this identity to compute 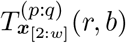 based on other terms

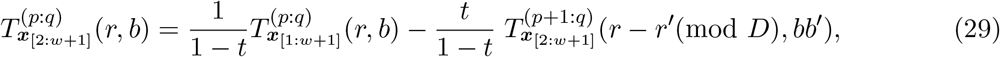

ov

Defining probability vector 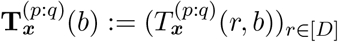, and calculating 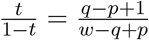, we can write the recursion in the vector form more concisely as:

♢ *Recursive delete* (*p ≤ q*):

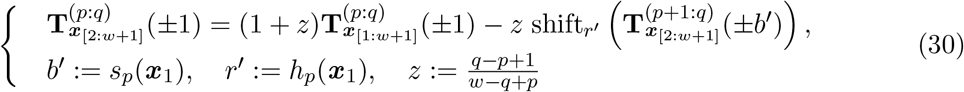

♢ *Recursive insert* (*p ≤ q*):

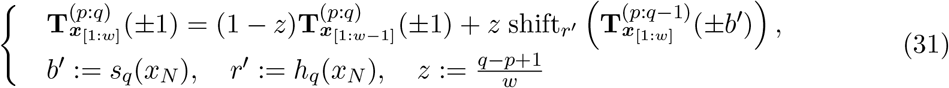

♢ *Basis (q < p*):

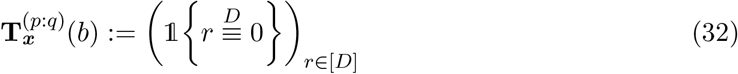

In order to implement an iterative algorithm, one can roll back the recursive calls and execute them in the reverse order. Namely for deletion, recursive relation for 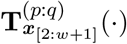 depends on 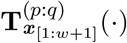, and 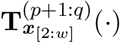, implying that updates for 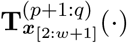 must precede updates for 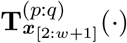, imposing a partial order corresponding to tuple length (*q — p* + 1). This in turn corresponds with the random subsequence length. For insertion recursion (18), sub-problems correspond to shorter subsequence lengths, while for deletion recursion (31), sub-problems correspond to longer subsequence lengths, which impose an ascending and descending dependency on subsequence length, respectively.

## Appendix B Parameter selection

The parameters for each model were selected according to the highest Spearman’s rank correlation. Figure S1 shows the accuracy of each method as a function of its parameters. The only model that was optimized for more than one parameter is OMH, where tuple length *t* and *k*-mer length *k* were jointly optimized.

### B.1 Hashing algorithm

In our parameter search both *CRC32* and *MurmurHash3* hashing algorithms were used to investigate the effect of hashing on the accuracy of min-hashing methods. We did not observe a significant effect of hashing on the accuracy of min-hashing. Therefore, MurmurHash3 was used in all experiments following the grid search, as it is more commonly used in bioinformatics practice.

### B.2 Tensor Slide Sketch

As TSS produces a 2-dimensional output, in all experiments the embedding dimension was set to 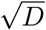. For example, if MH, WMH, OMH, and TS were run with 64 embedding dimensions, TSS sketch dimension was set to 8. TSS concatenates these 8-dimensional sketches every s steps. Throughout the experiments the length of window and stride were always set proportional to sequence length, 10% and 1% of the length respectively. As a result, the overall sketch size of TSS remains comparable to other methods.

**Fig. S1:**
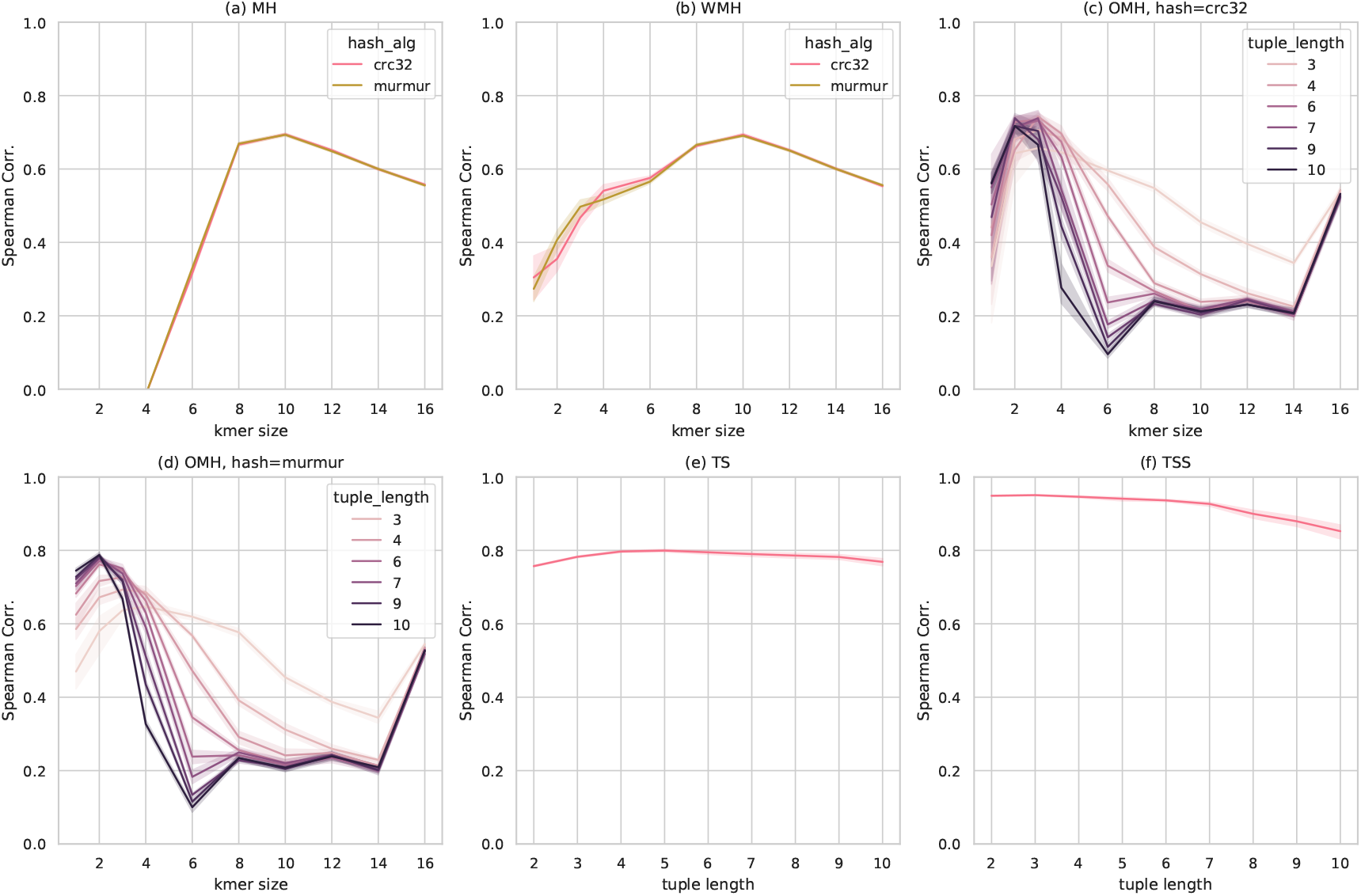
Hyper-parameters for each method: subplots (a) to (f) show the Spearman’s rank correlation of each sketching method versus their respective parameters. The shades in the plots indicate the standard deviation of Spearman’s correlation, calculated for over independent runs of the same configuration. The grid search is defined over 1 ≤ *k* ≤ 16 and 2 ≤ *t* ≤ 10. The tuple length *t* = 1 is not included, because such a case is covered by WMH. In subplot (a) the accuracy of MH is depicted versus the *k*-mer size, and the hashing algorithm is indicated by the color. (b) similar to subplot (a), but for WMH, In subplot (c), Spearman’s correlation of OMH is for *k*-mer size and tuple length, with the hashing algorithm fixed to CRC32. Subplot (d) is similar to (c), and the only difference is that the hashing method is set to MurmurHash3. Subplots (e) and (f) illustrate the average accuracy of TS and TSS respectively, as a function of tuple length *t*.

## Footnotes

4 The plus-minus signs are shorthand for two equations: one by taking all to be +, and another taking all to be —, but not any other combination, for example sin(*a ± b*) = sin(*a*) cos(*b*) ± sin(*b*) cos(*a*). If there are ∓ in the equation, their order for execution must the reversed, for example cos(*a ± b*) = cos(*a*) cos(*b*) ∓ sin(*a*) sin(*b*).

## References

1. Apostolico, A., Denas, O.: Efficient tools for comparative substring analysis. Journal of Biotechnology 149(3), 120–126 (2010). DOI 10.1016/J.JBIOTEC.2010.05.006. URL https://www.sciencedirect.com/science/article/pii/S0168165610002403?via73Dihub

2. Broder, A.Z.: On the resemblance and containment of documents. Proceedings of the International Conference on Compression and Complexity of Sequences pp. 21–29 (1997)

3. Hahn, L., Leimeister, C.A., Ounit, R., Lonardi, S., Morgenstern, B.: rasbhari: Optimizing Spaced Seeds for Database Searching, Read Mapping and Alignment-Free Sequence Comparison. PLoS computational biology 12(10), e1005, 107 (2016). DOI 10.1371/journal.pcbi.1005107. URL http://www.ncbi.nlm.nih.gov/pubmed/27760124 http://www.pubmedcentral.nih.gov/articlerender.fcgi?artid=PMC5070788

4. Kulis, B., Grauman, K.: Kernelized locality-sensitive hashing for scalable image search. In: 2009 IEEE 12th international conference on computer vision, pp. 2130–2137. IEEE (2009)

5. Leimeister, C.A., Boden, M., Horwege, S., Lindner, S., Morgenstern, B.: Fast alignment-free sequence comparison using spaced-word frequencies. Bioinformatics (Oxford, England) 30(14), 1991–9 (2014). DOI 10.1093/bioinformatics/btu177. URL http://www.ncbi.nlm.nih.gov/pubmed/24700317 http://www.pubmedcentral.nih.gov/articlerender.fcgi?artid=PMC4080745

6. Leimeister, C.A., Morgenstern, B.: Kmacs: the k-mismatch average common substring approach to alignment-free sequence comparison. Bioinformatics (Oxford, England) 30(14), 2000–8 (2014). DOI 10.1093/bioinformatics/btu331. URL http://www.ncbi.nlm.nih.gov/pubmed/24828656 http://www.pubmedcentral.nih.gov/articlerender.fcgi?artid=PMC4080746

7. Ma, B., Tromp, J., Li, M.: PatternHunter: faster and more sensitive homology search. Bioinformatics 18(3), 440–445 (2002). DOI 10.1093/bioinformatics/18.3.440. URL http://www.ncbi.nlm.nih.gov/pubmed/11934743 https://academic.oup.com/bioinformatics/article-lookup/doi/10.1093/bioinformatics/18.3.440

8. Mak, D.Y., Benson, G.: All hits all the time: parameter-free calculation of spaced seed sensitivity. Bioinformatics 25(3), 302–308 (2009). DOI 10.1093/bioinformatics/btn643. URL https://academic.oup.com/bioinformatics/article-lookup/doi/10.1093/bioinformatics/btn643

9. Marcais, G., Deblasio, D., Pandey, P., Kingsford, C.: Locality-sensitive hashing for the edit distance. Bioinformatics 35(14), i127–i135 (2019). DOI 10.1093/bioinformatics/btz354

10. Noé, L.: Best hits of 11110110111: model-free selection and parameter-free sensitivity calculation of spaced seeds. Algorithms for molecular biology: AMB 12, 1 (2017). DOI 10.1186/s13015-017-0092-1. URL http://www.ncbi.nlm.nih.gov/pubmed/28289437 http://www.pubmedcentral.nih.gov/articlerender.fcgi?artid=PMC5310094

11. Noe, L., Martin, D.E.: A coverage criterion for spaced seeds and its applications to support vector machine string kernels and k-mer distances. Journal of Computational Biology 21(12), 947–963 (2014). DOI 10.1089/cmb.2014.0173. URL http://www.ncbi.nlm.nih.gov/pubmed/25393923 http://www.pubmedcentral.nih.gov/articlerender.fcgi?artid=PMC4253314

12. Ondov, B.D., Treangen, T.J., Melsted, P., Mallonee, A.B., Bergman, N.H., Koren, S., Phillippy, A.M.: Mash: Fast genome and metagenome distance estimation using MinHash. Genome Biology 17(1), 1–14 (2016). DOI 10.1186/s13059-016-0997-x. URL http://dx.doi.org/10.1186/s13059-016-0997-x

13. Onodera, T., Shibuya, T.: The Gapped Spectrum Kernel for Support Vector Machines. In: International Workshop on Machine Learning and Data Mining in Pattern Recognition, pp. 1–15. Springer, Berlin, Heidelberg (2013). DOI 10.1007/978-3-642-39712-7\_1. URL http://link.springer.com/10.1007/978-3-642-39712-7_1

14. Pham, N., Pagh, R.: Fast and scalable polynomial kernels via explicit feature maps. In: Proceedings of the 19th ACM SIGKDD international conference on Knowledge discovery and data mining, pp. 239–247 (2013)

15. Rohling, S., Dencker, T., Morgenstern, B.: The number of k-mer matches between two DNA sequences as a function of k. bioRxiv p. 527515 (2019). DOI 10.1101/527515. URL https://www.biorxiv.org/content/10.1101/527515v2

16. Sims, G.E., Jun, S.R., Wu, G.A., Kim, S.H.: Whole-genome phylogeny of mammals: Evolutionary information in genic and nongenic regions. Proceedings of the National Academy of Sciences of the United States of America 106(40), 17, 077-17,082 (2009). DOI 10.1073/pnas.0909377106. URL http://www.ncbi.nlm.nih.gov/pubmed/19805074 http://www.pubmedcentral.nih.gov/articlerender.fcgi?artid=PMC2761373

17. Sims, G.E., Kim, S.H.: Whole-genome phylogeny of Escherichia coli/Shigella group by feature frequency profiles (FFPs). Proceedings of the National Academy of Sciences of the United States of America 108(20), 8329–34 (2011). DOI 10.1073/pnas.1105168108. URL http://www.ncbi.nlm.nih.gov/pubmed/21536867 http://www.pubmedcentral.nih.gov/articlerender.fcgi?artid=PMC3100984

18. Ulitsky, I., Burstein, D., Tuller, T., Chor, B.: The Average Common Substring Approach to Phylogenomic Reconstruction. Journal of Computational Biology 13(2), 336–350 (2006). DOI 10.1089/cmb.2006.13.336. URL http://www.liebertpub.com/doi/10.1089/cmb.2006.13.336

